# Depolarizing GABA Transmission Restrains Activity-Dependent Glutamatergic Synapse Formation in the Developing Hippocampal Circuit

**DOI:** 10.1101/742148

**Authors:** Christopher K. Salmon, Horia Pribiag, W. Todd Farmer, Scott Cameron, Emma V. Jones, Vivek Mahadevan, David Stellwagen, Melanie A. Woodin, Keith K. Murai

**Affiliations:** Centre for Research in Neuroscience, Department of Neurology and Neurosurgery, The Research Institute of the McGill University Health Centre, Montreal General Hospital, Montreal, Quebec, H3G 1A4, Canada; Department of Cell & Systems Biology, University of Toronto, Toronto, Ontario, M5S 3G5, Canada

**Keywords:** Synapse formation, hippocampus, GABA transmission, dendritic spines, chloride homeostasis, KCC2, circuit development, autism

## Abstract

GABA is the main inhibitory neurotransmitter in the mature brain but has the paradoxical property of depolarizing neurons during early development. Depolarization provided by GABA_A_ transmission during this early phase regulates neural stem cell proliferation, neural migration, neurite outgrowth, synapse formation, and circuit refinement, making GABA a key factor in neural circuit development. Importantly, depending on the context, depolarizing GABA_A_ transmission can either drive neural activity, or inhibit it through shunting inhibition. The varying roles of depolarizing GABA_A_ transmission during development, and its ability to both drive and inhibit neural activity, makes it a difficult developmental cue to study. This is particularly true in the later stages of development, when the majority of synapses form and GABA_A_ transmission switches from depolarizing to hyperpolarizing. Here we addressed the importance of depolarizing but inhibitory (or shunting) GABA_A_ transmission in glutamatergic synapse formation in hippocampal CA1 pyramidal neurons. We first showed that the developmental depolarizing-to-hyperpolarizing switch in GABA_A_ transmission is recapitulated in organotypic hippocampal slice cultures. Based on the expression profile of K^+^-Cl^-^ co-transporter 2 (KCC2) and changes in the GABA reversal potential, we pinpointed the timing of the switch from depolarizing to hyperpolarizing GABA_A_ transmission in CA1 neurons. We found that blocking depolarizing but shunting GABA_A_ transmission increased excitatory synapse number and strength, indicating that depolarizing GABA_A_ transmission can restrain glutamatergic synapse formation. The increase in glutamatergic synapses was activity-dependent, but independent of BDNF signalling. Importantly, the elevated number of synapses was stable for more than a week after GABA_A_ inhibitors were washed out. Together these findings point to the ability of immature GABAergic transmission to restrain glutamatergic synapse formation and suggest an unexpected role for depolarizing GABA_A_ transmission in shaping excitatory connectivity during neural circuit development.

## INTRODUCTION

γ-Aminobutyric acid (GABA) is the main inhibitory neurotransmitter in the mature brain. However, GABA is paradoxically depolarizing during nervous system development. Many *in vitro* studies in rodents have shown that depolarizing GABA_A_ transmission provides excitatory drive during gestation and early postnatal CNS development, driving early network oscillations (ENOs) thought to promote activity-dependent maturation of neural circuits (Ben-Ari et al., 2012). However, recent work suggests that despite providing local depolarization, immature GABA_A_ transmission has inhibitory effects *in vivo* (Kirmse et al., 2015; Oh et al., 2016; Valeeva et al., 2016). This ability of GABA to be simultaneously depolarizing and inhibitory relies on shunting inhibition, which results from a decrease in input resistance and membrane time constant when GABA_A_ receptors open, regardless of the direction of CI^-^ flux (Staley and Mody, 1992). Importantly, shunting inhibition can occur in conjunction with both hyperpolarizing and depolarizing GABA_A_ transmission, and we therefore refer to the latter case as depolarizing/inhibitory.

Depolarizing GABA_A_ transmission is implicated in numerous neurodevelopmental processes in vertebrates, including neural stem cell proliferation (Liu et al., 2005), cell migration (Behar et al., 2000), neurite outgrowth (Cancedda et al., 2007), synapse formation, and circuit refinement (Akerman and Cline, 2006; Cancedda et al., 2007; Wang and Kriegstein, 2008). Critically, circuit activity supported by depolarizing GABA_A_ transmission *in vitro* drives calcium influx thought to be important for glutamatergic synapse development (Leinekugel et al., 1995; Ben-ari et al., 1997; Griguoli and Cherubini, 2017). Indeed, disrupting the depolarizing nature of GABA_A_ transmission by interfering with chloride homeostasis alters glutamatergic synapse formation and maturation (Akerman and Cline, 2006; Wang and Kriegstein, 2008). However, the effects of GABA_A_ transmission itself on glutamatergic synapse development and the timing of these effects remain poorly defined. This is partly due to the difficulty in manipulating depolarizing GABA_A_ transmission in defined cell types and circuits with sufficient temporal resolution to specifically target the period when glutamatergic synapses are forming, while sparing the preceding developmental roles of GABA. Several studies have prematurely hyperpolarized the reversal potential for chloride (E_Cl_) by disrupting chloride homeostasis for more than a week during perinatal development, across a timespan in which the targeted neurons terminally divide, migrate, extend neurites and are incorporated into the surrounding circuitry (Ge et al., 2006; Cancedda et al., 2007; Wang and Kriegstein, 2008). This work suggests that disrupting E_Cl_ alters neurite and synapse maturation, however, it has been noted that additional studies with higher temporal resolution are needed (Akerman and Cline, 2007; Kirmse et al., 2018). Closing this gap in our understanding of how GABA_A_ transmission and its transition from a depolarizing to a hyperpolarizing state impacts glutamatergic synapse development will help solve a now classic problem in developmental neurobiology, and will likely be of clinical significance as disruptions of GABA_A_ transmission during brain development are associated with neurodevelopmental disorders (El Marroun et al., 2014; He et al., 2014; Tyzio et al., 2014).

Here we investigated the role of depolarizing GABA_A_ transmission in glutamatergic synapse formation on hippocampal CA1 pyramidal cells. To perform temporally precise pharmacological manipulations of GABA_A_ transmission during neural circuit development, we took advantage of the properties of the organotypic hippocampal slice culture. This preparation preserves the anatomy and the developmental progression of the hippocampus, including the time course of excitatory synapse formation (Buchs et al., 1993; Muller et al., 1993; De Simoni et al., 2003). This system enabled us to define a narrow time window during the first week of slice development in which GABA_A_ transmission shifts from immature, depolarizing transmission, to hyperpolarizing transmission in CA1 pyramidal cells. Previous work suggests that blocking depolarizing GABA_A_ transmission during development will remove excitatory drive and decrease excitatory synapse formation and maturation (Ben-Ari et al., 2007; Wang and Kriegstein, 2008). Contrary to these predictions, we found that transient blockade of immature, depolarizing GABA_A_ transmission increased glutamatergic synapse number and function on CA1 pyramidal cells. This unexpected effect was explained by the finding that, at this stage of development, depolarizing GABA_A_ transmission provides shunting inhibition, which when blocked alleviated a restraint on activity-dependent synapse formation. Interestingly, the activity-dependent increase in glutamatergic synapses was stable for at least a week. Furthermore, the effect could not be reproduced by prematurely hyperpolarizing E_GABA_, and was independent of BDNF signalling. Our results therefore point to an important time window during hippocampal development when immature GABA_A_ transmission can restrain excitatory synapse development, and that interfering with GABA_A_ transmission at this stage can have lasting effects on neural circuitry.

## RESULTS

### GABA_A_ transmission switches from depolarizing to hyperpolarizing in CA1 cells during the first week in hippocampal slice culture

Depolarizing GABA_A_ transmission relies on relatively high intracellular chloride ([Cl^-^]_i_). As neurons mature during the first weeks of postnatal CNS development, Na^+^-K^+^-Cl^-^ cotransporter (NKCC1) expression is downregulated and K^+^-Cl^-^ cotransporter 2 (KCC2) is upregulated, lowering [Cl^-^]_i_ (Rivera et al., 1999; Yamada et al., 2004). GABA_A_ receptors are largely permeable to Cl^-^, and to a lesser extent bicarbonate (HCO_3_^-^) (Kaila, 1994; Staley and Proctor, 1999). When [Cl^-^]_i_ lowers to the point at which the reversal potential for GABA (E_GABA_) hyperpolarizes below the resting membrane potential, GABA_A_ transmission switches from depolarizing to hyperpolarizing. To pinpoint when this switch from depolarization to hyperpolarization occurs in CA1 pyramidal cells in hippocampal organotypic slices, we first assessed the timing of KCC2 upregulation across the first two weeks *in vitro* and found expression of both KCC2 monomers (KCC2-M) and oligomers (KCC2-O) underwent a large and graded increase between 3 and 7DIV (Fig 1A,B), reaching near-maximal levels by 7 days *in vitro* (DIV) (Fig 1B). Using this timeframe as a guide, we performed gramicidin perforated patch recordings to determine the GABA_A_ reversal potential (E_GABA_) in CA1 pyramidal cells (exemplary traces and IV curves shown in Figures 1C and D). At 3-4 DIV, E_GABA_ was depolarized with respect to resting membrane potential (RMP) (Fig 1E-G). However, by 6-7 DIV E_GABA_ was hyperpolarized with respect to RMP, indicating a switch to hyperpolarizing GABA_A_ transmission by 6-7 DIV (Fig 1C-G), a timeframe similar to that reported previously for CA1 pyramidal cells (Swann et al., 1989). E_GABA_ was more negative than action potential threshold at 3-4 DIV (Fig 1E,G), suggesting GABA is depolarizing but not capable of directly depolarizing neurons past action potential (AP) threshold from rest at this stage.

**Figure 1.**
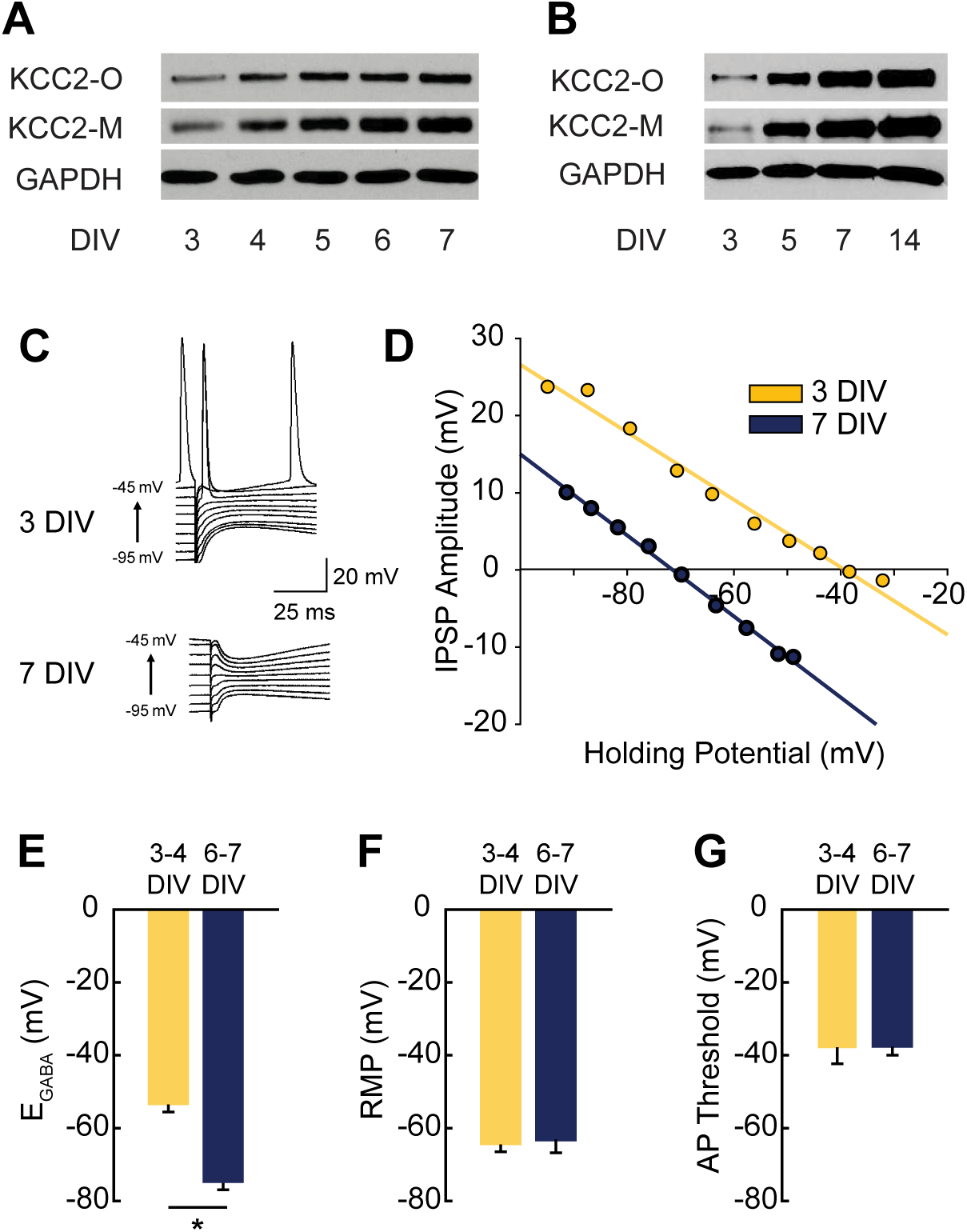
GABA reversal potential (E_GABA_) shifts from depolarizing to hyperpolarizing between 3 and 7 DIV. ***A-B***, High (A) and low (B) temporal resolution western blots showing increasing expression of KCC2 monomers (KCC2-M) and oligomers (KCC2-O). ***C-D***, Representative traces and representative IV curves from GABAergic responses at 3DIV and 7DIV. ***E***, E_GABA_ summary plots (3/4 DIV: −53.3±6.1mV, n=5; 6/7 DIV: −74.7± 6.4mV, n=5, p=0.04). ***F,*** Resting membrane potential summary plots (3/4 DIV: −64.5±2.3mV, n=5; 6/7 DIV: −63.4 ± 3.8mV, n=5). ***G***, Action potential threshold summary plot (3/4 DIV: −38.2 ± 4.2mV, n=5; 6/7 DIV-37.7 ± 2.3mV, n=5).

### Blocking depolarizing GABA_A_ transmission increases CA1 spine density

Overexciting mature neurons by blocking hyperpolarizing GABA_A_ transmission is known to cause a collapse of dendritic spines both *in vivo* (Zeng et al., 2007) and *in vitro* (Muller et al., 1993; Drakew et al., 1996; Jourdain et al., 2002; Zha et al., 2005). In particular, applying GABA_A_ antagonists to organotypic hippocampal cultures at 5 or 23 DIV over a period of 2 to 3 days was shown to cause a robust loss of spines (Drakew et al., 1996; Zha et al., 2005). Consistent with this, when we blocked GABA_A_ transmission with the GABA_A_R antagonist, bicuculline (BIC) from 5-7 DIV (when GABA_A_ transmission is hyperpolarizing (Fig 1C-H)), spine density decreased by 34% (Fig 2A-C). This suggests that by this stage, excitatory transmission causes overexcitation and spine loss in the absence of hyperpolarizing GABA_A_ transmission.

**Figure 2.**
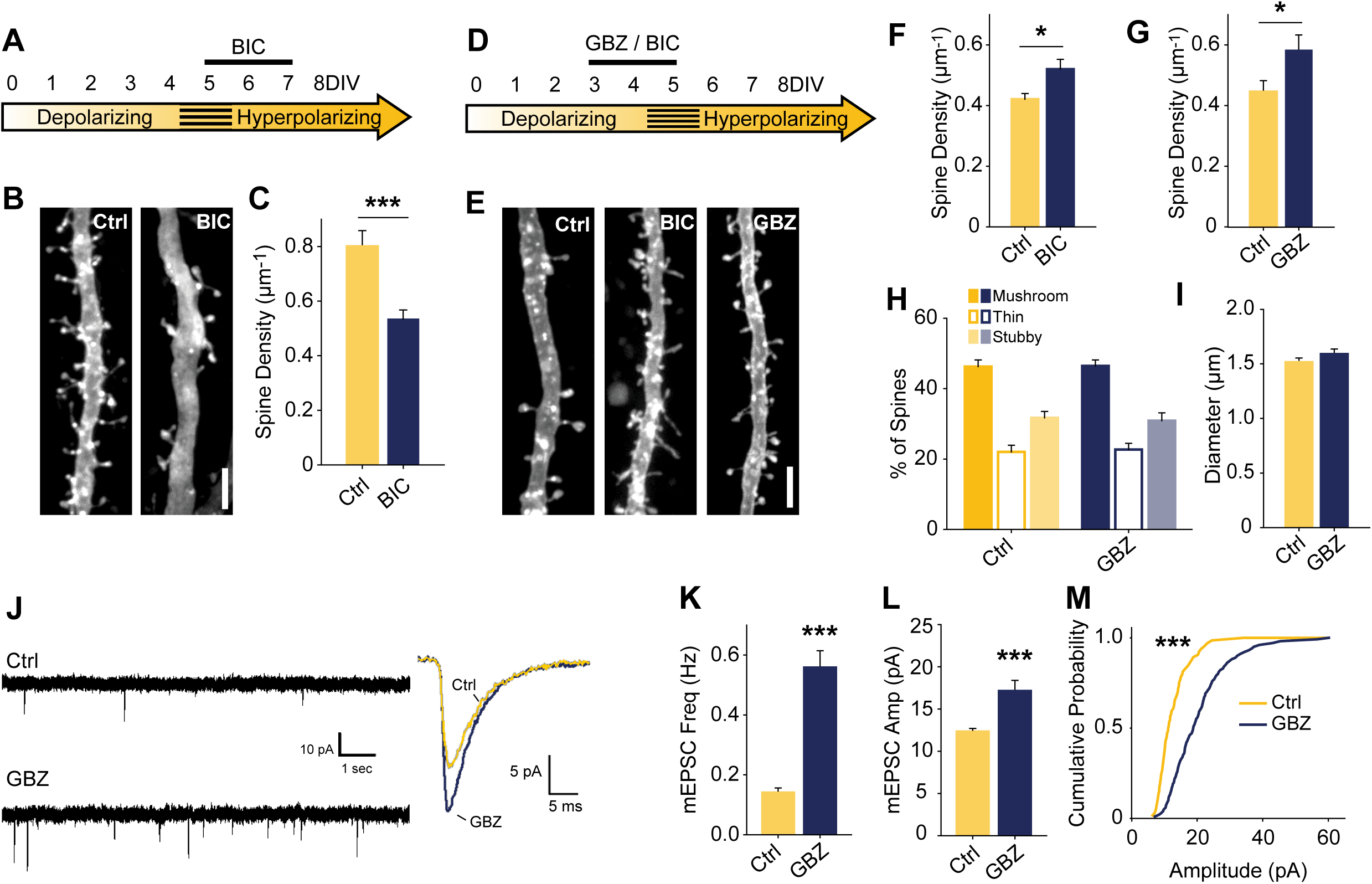
Blocking depolarizing GABA_A_ transmission increases excitatory synapse number. ***A***, Time course of bicuculline (BIC) treatment for B-C. ***B-C***, Spine density after 5-7 DIV BIC treatment (Control 0.80±0.06 spines/um, n=36, BIC 0.53±0.03, n=50; N=3; p<0.001, Mann-Whitney).***D***, Time course of pharmacological treatments for E-M. ***E-G***, Spine density after 3-5 DIV GBZ (G: Control 0.44±0.12 spines/um, n=145, GBZ 0.58±0.17, n=77; N=11; p=0.04) and BIC treatment (F: Control 0.42±0.02 spines/um, n=55, BIC 0.52±0.03 spines/um, n=41; N=9; P=0.027, Mann-Whitney). ***H,I***, 3D spine morphology and dendrite diameter after GBZ. ***J***, Representative traces of mEPSCs. ***K***, mEPSC frequency summary plot (Control 0.14±0.02 Hz, GBZ 0.56±0.06 Hz, p<0.001, Mann-Whitney). ***L***, mEPSC amplitude summary plot (Control 12.32±0.37 pA, n=8, GBZ 17.12±1.27 pA, n=10, p<0.001, Mann-Whitney). ***M***, Cumulative distributions of amplitudes (p<0.0001, Kolmogorov-Smirnov test). Scale bars 3µm.

To assess the role of immature, depolarizing GABA_A_ transmission on dendritic spine development, we inhibited GABA_A_ transmission earlier, from 3-5 DIV (Fig 2D). Previous work suggests that inhibiting depolarizing GABA_A_ transmission during development would decrease glutamatergic synapse formation and maturation (Ben-ari et al., 1997; Hanse et al., 1997; Cancedda et al., 2007; Wang and Kriegstein, 2008). However, in contrast to these findings, BIC applied for 48 hours from 3 to 5 DIV significantly increased dendritic spine density (25% increase)(Fig 2E-F). This effect was fully reproducible with the GABA_A_R antagonist gabazine (GBZ)(31% increase)(Fig 2E,G), which is a more specific antagonist of GABA_A_Rs (Heaulme et al., 1986) and blocks inhibition more consistently in hippocampal neurons (Sokal et al., 2000). We also verified that the presence of penicillin-streptomycin in the culture medium was not associated with this effect by blocking GABA transmission in the absence antibiotics, and found the same increase in dendritic spines (S1A-C Fig).

To assess whether the supernumerary spines induced by blocking depolarizing GABA_A_ transmission showed structural differences, we analyzed spine morphology. GBZ treatment did not affect the proportions of mushroom, thin, and stubby spines (Fig 2H), 2-dimensional head area (Control: 0.32±0.02 µm^2^; GBZ: 0.37±0.04 µm^2^, p>0.10), head diameter (Control: 0.58±0.02 µm^2^; GBZ: 0.62±0.03 µm^2^, p>0.1), spine length (Control 1.66±0.09 µm^2^; GBZ: 1.83±0.08 µm^2^, p>0.1) or dendrite diameter (Fig 2I).

We next asked whether the increased number of spines constituted an increase in *bona fide* glutamatergic synapses on CA1 cells by recording miniature EPSCs (mEPSC). Consistent with the increase in dendritic spine density, mEPSC analysis showed that GBZ treatment (3-5 DIV) increased mESPC frequency 3-fold (Fig 2J,K). Miniature EPSC amplitude also increased, indicating enhanced synaptic strength (Fig 2L-M). Together, these results suggest that immature GABA_A_ transmission restrains glutamatergic synapse formation and maturation.

The narrow time window we examined raised the possibility that the spine-enhancing effect of GABA_A_ blockade is limited to a short period directly prior to the depolarizing to hyperpolarizing shift in GABA_A_ transmission. This would suggest that GABA_A_ transmission restrains glutamatergic synapse formation only during a very short transition state. To test whether this was the case, we prepared slices 3 days earlier (P2) and applied GBZ at 3DIV for 48h (S1D Fig). We found that GABA_A_R blockade in these younger slices also caused a significant increase in spines (S1E,F Fig), suggesting that depolarizing GABA_A_ transmission is capable of restraining synapse formation for an appreciable period during postnatal development.

### Bumetanide treatment has no effect on spine numbers

Previous work suggests that abrogating GABAergic depolarization by prematurely rendering GABA hyperpolarizing decreases glutamatergic synapse formation (Ge et al., 2006; Wang and Kriegstein, 2008). However, our data show that a complete loss of depolarizing GABA_A_ transmission increases glutamatergic synapse formation. These contrasting results raise the question of whether the depolarizing nature of GABA_A_ transmission is important for the normal development of glutamatergic synapse number in our period of interest (3-5 DIV). To address this, we asked whether prematurely hyperpolarizing E_GABA_ could mimic the effect of GABA_A_ blockade by treating slices with the NKCC1 blocker bumetanide (BUME) from 3 to 5DIV (S2 Fig). BUME is well established to lower E_GABA_ in immature neurons (Dzhala et al., 2005) and prematurely render GABA hyperpolarizing (Wang and Kriegstein, 2011). Treating slice cultures at 3DIV with BUME did not alter spine density on its own (S2A,B Fig), indicating that the depolarizing nature of GABA is not important for regulating spine numbers at this stage of development. Furthermore, BUME did not alter the effect of GBZ on spine density, indicating that the extent to which E_GABA_ is depolarized is not important for limiting spine density to normal levels at this stage.

Since KCC2 overexpression can cause an increase in spines through its non-transport, scaffolding function (Li et al., 2007; Fiumelli et al., 2012), we also assessed KCC2 expression following GBZ treatment. GBZ did not significantly elevate expression of KCC2 oligomers or monomers (S2C-E Fig).

### Driving depolarizing GABA_A_ transmission does not alter glutamatergic synapse number

Next, we investigated if increasing GABA_A_ transmission over the 3-5 DIV period would have the opposite effect to GABA-blockade and reduce excitatory synapses. Previous work has demonstrated that propofol, a positive allosteric modulator of GABA_A_Rs, decreases spine density in developing layer 2/3 principal cells of the somatosensory cortex when administered to rat pups over a 6h period at postnatal day 10, when GABA_A_ transmission is still depolarizing (Puskarjov et al., 2017). To test this in CA1 pyramidal cells, we increased depolarizing GABA_A_ transmission by administering muscimol (MUS) or diazepam (DZP) from 3 to 5DIV. MUS treatment did not significantly decrease spine density (S3A-C Fig). Furthermore, mEPSC frequency was unchanged, confirming MUS did not alter synapse numbers (S3D,E Fig). MUS has varying effects on different GABA receptors and can cause GABA_A_ receptor desensitization, making its effects difficult to interpret (Heck et al., 2007; Mortensen et al., 2010; Johnston, 2014). We therefore also tested whether enhancing GABA_A_ transmission with DZP could decrease glutamatergic synapses, but this also had no effect on spine density or mEPSCs (S3A-F Fig). Based on these results, increasing GABA_A_ transmission was not sufficient to decrease glutamatergic synapse number or function, suggesting depolarizing GABA_A_ transmission can only limit synapse formation up to a certain point at this stage of development. However, our results do not rule out the possibility that enhancing immature GABA_A_ transmission on different timescales or in other systems decreases glutamatergic synapse formation (Puskarjov et al., 2017).

### Increased glutamatergic synapses following blockade of depolarizing GABA_A_ transmission is activity-dependent

Based on our recordings showing that at 3-4DIV E_GABA_ is depolarized relative to RMP, but lower than action potential threshold (Fig 1E-G), GABA is likely to mediate shunting inhibition (ie depolarizing/inhibitory transmission), at this stage (see S4A Fig for schematic). To test this possibility, we puffed GABA locally while recording spontaneous or electrically evoked firing. GABA inhibited both spontaneous (Fig 3A,B) and evoked spiking (S1B,C Fig), suggesting that although E_GABA_ is depolarizing relative to RMP, GABA_A_ transmission is inhibitory during the 3-5 DIV timeframe. Blocking this depolarizing/inhibitory GABA_A_ transmission likely increased activity in our preparation, suggesting that the increase in glutamatergic synapses following GABA-blockade at 3DIV may be driven by activity-dependent mechanisms (Balkowiec and Katz, 2002; Pérez-Gómez and Tasker, 2013). To address this hypothesis, we measured levels of *Bdnf* and *Fos* mRNA, two activity regulated genes associated with glutamatergic synapse formation (Vicario-Abejón et al., 1998, 2002; Tyler and Pozzo-Miller, 2003; Chapleau et al., 2009). Both transcripts were significantly upregulated following 48-hour blockade of depolarizing/inhibitory GABA_A_ transmission (*Bdnf*: 5-fold increase, *Fos*: 2.5-fold increase) (Fig 3C). GABA_A_ blockade at 3DIV also increased Fos protein expression after 24 and 48 hours (Fig 3D-E). Together these data indicate that blocking immature depolarizing GABA_A_ transmission at this point caused an increase in activity in CA1 pyramidal cells. To test whether the increased synapse formation we observed following GABA blockade at 3DIV was activity-dependent, we treated slice cultures with GBZ and/or TTX, and found that while TTX alone had no effect on spine density, TTX blocked the GBZ-induced increase in spines (Fig 3F). From this we conclude that depolarizing/inhibitory GABA_A_ limits activity-dependent glutamatergic synapse formation at this point in the development of hippocampal circuitry in slice culture.

**Figure 3.**
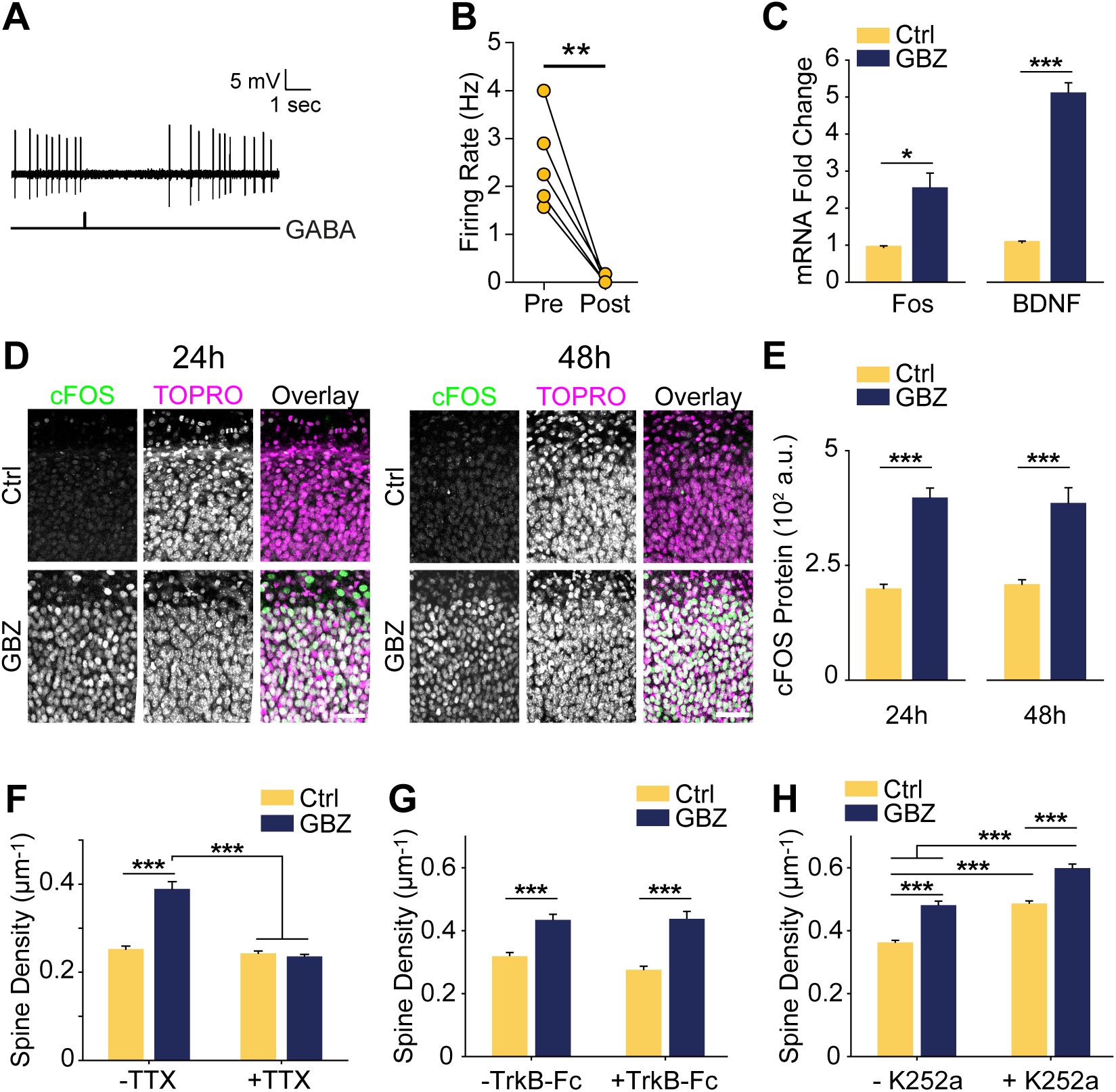
Increase in spine density following blockade of depolarizing/inhibitory GABA_A_ transmission is activity-dependent but does not rely on BDNF signalling. *A,* Sample trace of spontaneous activity inhibited by puffing on GABA. The line trace below indicates time of GABA puff. *B,* Summary plots of spontaneous activity pre- and post-GABA puff. *C*, BDNF and Fos transcript levels following GBZ from 3-5DIV (BDNF: Ctrl 1.07±0.04, GBZ 5.08±0.3, N=3, p<0.001; Fos: Ctrl 0.94±0.04, GBZ 2.52±0.4, N=3, p=0.02). *D,* Fos immunofluorescence 24 and 48 hours after GBZ treatment at 3DIV. Images depict the dividing line between stratum oriens (upper portion of panels) and stratum pyramidale (lower portion of panels) in area CA1. TOPRO-3-Iodide was used to visualize nuclei. *E*, Quantification of Fos immunofluorescence following 24h (Ctrl 196.5±12.2 au, n=10, GBZ 394.4±24.0 au, n=11, p<0.001) and 48h (Ctrl 205.6±12.1 au, n=10, GBZ 382.7±36.3 au, n=10, p<0.001, Mann-Whitney) of GBZ treatment which started at 3DIV. **F,** Quantification of spine density following GBZ and/or TTX treatment beginning at 3DIV (Ctrl 0.25±0.01 μm^-1^, n= 196, GBZ 0.39±0.01 μm^-1^, n=110, TTX 0.24±0.01 μm^-1^, n= 166, GBZ+TTX 0.23±0.01μm^-1^, n=154; N=5). Two-way ANOVA indicates a significant interaction between GBZ and TTX conditions, p<0.001. Significant differences between GBZ and all other conditions, p<0.001, Tukey post test. *G*, Quantification of spine density following GBZ and/or TrkB-Fc treatment (Ctrl 0.31±0.02, n=86, GBZ 0.42±0.02, n=68, TrkB-Fc 0.27±0.02, n=96, TrkB-Fc+GBZ 0.43±0.02, n=61; N=3; 2 Way ANOVA, no interaction, Tukey post test). *H*, Quantification of spine density following GBZ and/or K252a treatment (Ctrl 0.35±0.01, n=198, GBZ 0.49±0.03, n=144, K252a 0.47±0.02, n= 216, K252a+GBZ 0.58±0.04, n=185; all significant differences <0.001, 2 Way ANOVA, no interaction, Tukey post test).

BDNF is known to regulate activity-dependent synapse formation and plasticity (Park and Poo, 2013), and we therefore asked whether BDNF signaling was responsible for the increase in spines following blockade of depolarizing/inhibitory GABA_A_ transmission. We inhibited BDNF signalling during the 3 to 5DIV GBZ treatment using TrkB-Fc bodies or K252a (Ji et al., 2010; Puskarjov et al., 2014), however neither approach blocked the increase in spine density (Fig 3G,H), suggesting that BDNF signalling is not necessary for the observed increase in spines.

### Blocking depolarizing GABA_A_ transmission leads to a sustained increase in glutamatergic synapses

The observed increase in spine density induced by blocking depolarizing/inhibitory GABA_A_ transmission may only lead to a transient alteration without a longer lasting effect on glutamatergic synapses. To determine whether blockade of GABA_A_ transmission caused a temporary or sustained increase in glutamatergic synapses, we treated slices with GBZ from 3-5 DIV and allowed them to recover for an additional 5-9 days in the absence of GBZ (Fig 4A). This temporary GABA_A_ blockade resulted in a 37% increase in spine density after a 5-day recovery period (Fig 4B,C). Furthermore, after this recovery period, CA1 cells had more thin spines than mushroom spines, a difference not present in the control condition (Fig 4D). No changes in dendrite diameter were observed (Fig 4E). To determine if transient GBZ treatment led to long-term functional changes in glutamatergic synapses, we recorded mEPSC frequency and amplitude after 8-9 days of recovery. We found that mEPSC frequency was enhanced by 79%, while mEPSC amplitude was unchanged at this stage (Fig 4F-I). Together these data suggest that inhibiting depolarizing GABA_A_ transmission during a narrow time window can lead to persistent changes in glutamatergic synapse number in the hippocampus.

**Figure 4.**
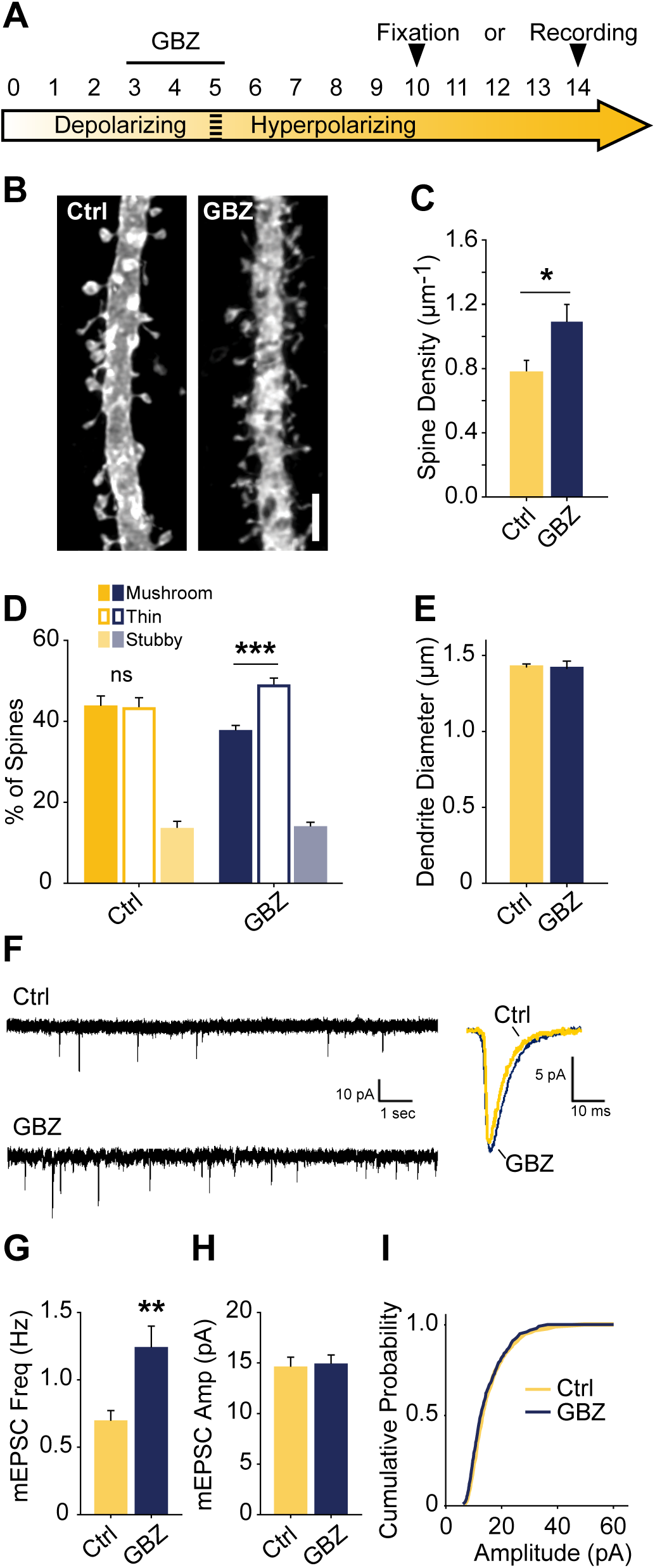
Transient blockade of depolarizing GABA_A_ transmission causes a lasting increase in excitatory synapses and alters spine morphology. *A,* Schematic time course of GBZ treatment and experimental endpoints. *B,C,* Spine density after 3-5 DIV GBZ treatment and 5 days of recovery (Control 0.78±0.08 spines/µm, n=127; GBZ washout 1.07±0.07 spines/µm, n=112; N=6; p=0.024,). *D,* 3D spine morphology after 5 days of recovery (***p<0.001, critical level 0.05, Two Way ANOVA with Holm Sidak Post Test). *E,* Dendrite diameter after recovery (p=0.86). *F,* Representative mEPSC traces from slices after 8-9 days recovery. *G,* mEPSC frequency summary plot (Control: 0.70±0.08 Hz, n=10 GBZ: 1.23±0.17 Hz, n=10, p=0.009). *H*, mEPSC amplitude summary plot (Control: 14.50±1.07 pA, n=10, GBZ: 14.80±1.00 pA, n=10, p=0.84). *I,* Cumulative mEPSC distributions (p=0.58, Kolmogorov-Smirnov test). Scale bar 3µm.

## DISCUSSION

Immature, depolarizing GABA_A_ transmission is believed to promote glutamatergic synapse formation and maturation (Ben-ari et al., 1997; Hanse et al., 1997; Wang and Kriegstein, 2009; Chancey et al., 2013). However, when and how GABA affects glutamatergic synapse formation remains to be fully understood. Indeed, several groups have noted that tools and approaches for manipulating depolarizing GABA_A_ transmission with higher temporal and spatial precision are needed to resolve this (Akerman and Cline, 2007; Chancey et al., 2013; Kirmse et al., 2018). We therefore sought to address the role of GABA_A_ transmission in glutamatergic synapse formation by performing precisely timed pharmacological manipulations in hippocampal slice cultures. We first mapped the depolarizing-to-hyperpolarizing shift of GABA_A_ transmission in CA1 cells. This was followed by structural and electrophysiological analysis which showed that blocking immature, depolarizing/inhibitory GABA_A_ transmission enhanced glutamatergic synapse function and number. Interestingly, the enhanced synapse number was stable following a recovery period. These results suggest that immature GABA_A_ transmission restrains glutamatergic synapse formation during an early phase of hippocampal circuit development. Using slice cultures allowed for more temporally precise manipulations that revealed this effect, though limitations of this model system must be considered when interpreting our results. In particular, exuberant glutamatergic synapse formation has been observed in slice cultures, and has been attributed to increases in distal dendritic branching (De Simoni et al., 2003). However, we minimized this confound by focusing on primary apical dendrites, which are fully formed by the time of pharmacological treatment. Thus, while further work will be required to extend our findings to other systems, the results of this study show that immature, depolarizing GABA_A_ transmission is capable of restraining glutamatergic synapse formation in certain contexts, and that the removal of this restraint by interfering with GABA_A_ transmission during development may cause a long-term increase in glutamatergic synapses.

### An unpredicted role for immature GABA_A_ transmission in restraining glutamatergic synapse formation

In the time window we examined, GABA_A_ transmission provides subthreshold depolarization and shunting inhibition, which when blocked alleviates a brake on glutamatergic synapse development. Taken in the context of previous work, our results suggest a couple of models for how immature GABA_A_ transmission affects hippocampal excitatory connectivity (S5 Fig). Firstly, the GABA-mediated restraint on glutamatergic synapse formation may be a short-lived feature of a depolarizing/inhibitory transition state that GABA passes through as E_Cl_ matures from depolarizing and excitatory to hyperpolarizing (Model 1, S5A-C Fig). However, recent work suggests GABA may be inhibitory throughout most or all of postnatal development. Therefore, in a second model, depolarizing but inhibitory GABA_A_ transmission may inhibit circuit activity from birth onward (Model 2, S5B-C Fig), thus restraining glutamatergic synapse formation across development.

The first model is based on evidence from acute slices suggesting that immature GABA_A_ transmission is capable of driving excitation (Gulledge and Stuart, 2003) and that depolarizing GABA_A_ transmission drives ENOs, which in turn promote glutamatergic synapse formation and unsilencing, and circuit refinement (Hanse et al., 1997; Ben-Ari, 2002; Wang and Kriegstein, 2009; Griguoli and Cherubini, 2017). Disrupting E_Cl_ or GABA_A_ transmission in this phase of development is hypothesized to interfere with synapse formation (S5A Fig), and this has been borne out by experimentally lowering E_Cl_ across the postmitotic period in immature neurons (Ge et al., 2006; Cancedda et al., 2007; Wang and Kriegstein, 2008). Incorporating our results refines this model and accounts for the role of GABA_A_ transmission in circuit development as it transitions from a depolarizing and excitatory to a hyperpolarizing state. Our work suggests that following an initial depolarizing phase in which GABA promotes excitation, as E_Cl_ progressively matures, GABA_A_ transmission passes through a transient but developmentally relevant depolarizing/inhibitory phase (S5B Fig). Such a transition phase is hinted at in the literature, as certain studies have shown that blocking depolarizing GABA_A_ transmission can silence ENOs, while others show GABA_A_ blockade to increase circuit activity by eliciting interictal discharges or paroxysmal activity (Le Magueresse et al., 2006; Ben-Ari et al., 2007). Our results suggest that during this transition phase, depolarizing GABA_A_ transmission is inhibitory and restrains glutamatergic synapse formation. Blocking GABA_A_ transmission at this time alleviates the restraint, allowing for activity-dependent synapse formation (S5B Fig). Following this transition phase, GABA_A_ transmission becomes fully hyperpolarizing as the glutamatergic system becomes capable of overexcitation. The result of GABA_A_ blockade at this stage is loss of spines (Fig 2 and S5C Fig) (Swann et al., 1989; Drakew et al., 1996; Zeng et al., 2007). Interestingly, the absence of a similar spine loss following blockade of depolarizing/inhibitory GABA_A_ transmission at 3DIV suggests that, while this still immature GABAergic inhibition is important for regulating activity levels, the glutamatergic system is not yet mature enough to cause a pathological collapse of synapse numbers similar to that seen in models of epilepsy (29,30).

Alternatively, in a second model, it is possible that depolarizing GABA_A_ transmission provides shunting inhibition throughout the postnatal period, thereby restraining synapse formation and circuit activity during development (S5B-C Fig, green shaded area). Indeed, emerging evidence suggests that depolarizing GABA_A_ transmission exerts inhibitory effects on ENOs *in vivo*, from at least P3 onward (Kirmse et al., 2015; Valeeva et al., 2016; Che et al., 2018). Consistent with this, our results in slices cultured from younger mice (S1D-F Fig) suggest that GABA_A_ transmission restrains synapse formation over a period of up to 5 days of hippocampal circuit development. While previous work has admittedly demonstrated that prematurely rendering GABA_A_ transmission hyperpolarizing *in vivo* decreases glutamatergic synapse formation (Ge et al., 2006; Cancedda et al., 2007; Wang and Kriegstein, 2008, 2011), it is noteworthy that these earlier studies manipulated E_Cl_ over extended periods that spanned multiple phases of postmitotic neuronal development, including cell migration, axonal/dendritic growth, synapse formation and circuit refinement. Depolarizing GABA_A_ transmission is thought to play important roles in all of these processes (Owens and Kriegstein, 2002), and hence the observed effects of prematurely reducing E_Cl_ on synapses may be secondary to other alterations in neuronal and circuit development. Indeed, soma size and dendritic branching are altered when GABA is prematurely rendered hyperpolarizing over an extended time period (Cancedda et al., 2007; Wang and Kriegstein, 2008). More temporally precise manipulations of GABA_A_ transmission and E_Cl_ are therefore essential for clarifying the roles of GABA during critical phases of synapse formation *in vivo*. Interestingly, the finding that propofol administered to postnatal day 10 rats decreased spine number supports the notion that there is a developmental period *in vivo* during which immature GABA_A_ transmission restrains glutamatergic synapse formation (Puskarjov et al., 2017).

When considering these two models, it is important to note that an inhibitory effect of depolarizing GABA_A_ transmission does not preclude a role for GABA in driving ENOs, as it has been demonstrated that depolarizing chloride currents are only involved in the initial generation of GDPs in acute slices, after which they inhibit the continuation of the same GDPs (Khalilov et al., 2015). Thus, depolarizing GABA_A_ transmission may simultaneously generate ENOs, while also maintaining control of wider circuit activity, thereby limiting runaway glutamatergic synapse formation. These dichotomous effects of GABA may rely on where GABAergic inputs impinge on the post synaptic neuron. Gulledge and Steward (2003) showed that in young rats, puffing GABA on distal dendrites of Layer 5 pyramidal cells facilitated firing, while puffing GABA on the cell body inhibited firing. Thus, different GABAergic interneuron subtypes may be responsible for driving ENOs vs restraining glutamatergic synapse formation. Furthermore, despite the evidence suggesting GABA is inhibitory throughout most of postnatal development *in vivo*, it has been shown that high frequency uncaging or stimulated release of GABA onto dendrites of layer 2/3 pyramidal cells in the neocortex can elicit formation of glutamatergic and GABAergic synapses during development *in vivo* (Oh et al., 2016). Although it remains to be seen whether endogenous patterns of GABA release can have similar effects, it appears there may be a local trophic role for depolarizing GABA_A_ transmission, which may promote synapse formation even as its circuit-wide inhibitory effects restrain the same process as we have demonstrated. More work is needed to dissect the possible roles of GABA in local synapse formation and more global circuit development, and to understand how the role of GABA changes across development.

### Sustained Changes in Glutamatergic Synapses and Neurodevelopmental Disorders

Remarkably, we found that a transient blockade of depolarizing, inhibitory GABA_A_ transmission led to a sustained increase in both the number of glutamatergic synapses and the proportion of thin spines, indicating that transient manipulations of immature GABA_A_ transmission can profoundly alter hippocampal connectivity (Fig 4). Using slice cultures allowed for more temporally precise manipulations that revealed this effect, though it remains to be seen if the phenomenon persists *in vivo.* These questions are clinically relevant, as a role for GABA in restraining synapse formation may change how we understand and mitigate the effects of anticonvulsants, anaesthetics and drugs of abuse on neonatal, as well as fetal development, as GABA is believed to be depolarizing mainly in late gestation in humans (Vanhatalo et al., 2005; Sedmak et al., 2015). Furthermore, both the persistent increase in synapses and spines and the shift in spine morphologies we observed after recovery from transient GBZ treatment are reminiscent of “spinopathies” seen in intellectual disabilities including Fragile X syndrome and autism spectrum disorders (Lacey and Terplan, 1987; Irwin et al., 2000, 2001; Kaufmann and Moser, 2000; Fiala et al., 2002; Hutsler and Zhang, 2010). Numerous models of ASDs are associated with a delay in the depolarizing to hyperpolarizing shift in E_GABA_ (He et al., 2014; Tyzio et al., 2014; Leonzino et al., 2016). Such a delayed transition to hyperpolarizing E_GABA_ likely translates to a delay in the onset of adequate shunting inhibition when GABA is still depolarizing, which may increase glutamatergic synapse formation in a manner similar to that which we observed when blocking depolarizing/inhibitory GABA_A_ transmission. Furthermore, mutation of the β3 GABA_A_ receptor subunit, the expression of which peaks during development when GABA is depolarizing, has been observed in ASD (Menold et al., 2001; Buxbaum et al., 2002; Chen et al., 2014). The findings presented in the current study may provide a causal link between these mutations and the hyperconnectivity observed in ASDs. Thus, further investigation is required to understand if impairments of depolarizing/inhibitory GABA_A_ transmission contribute to the lasting alterations of spines and synapses in these conditions. Finally, the possibility that GABA bidirectionally controls synapse formation may yield novel clinical approaches for correcting synaptic deficits in neurodevelopmental disorders.

## MATERIALS AND METHODS

### Animals

Experiments were approved by the Montreal General Hospital Facility Animal Care Committee and followed guidelines of the Canadian Council on Animal Care. Male and female C57BL6 mice kept on a 12:12 light-dark cycle were used to prepare organoptypic cultures.

### Slice Preparation

Organotypic hippocampal slices were prepared as described previously (Haber et al., 2006). Briefly, hippocampi were extracted from postnatal day 5 mice and cut into 300µm slices with a McIllwain tissue chopper (Stoelting). Slices were cultured on semiporous tissue culture inserts (Millipore) that sat in culture medium composed of minimal essential medium (MEM) supplemented with Glutamax (Invitrogen, Cat. No. 42360032), 25% horse serum (Invitrogen, Cat. No. 26050088), 25% HBSS (Invitrogen, Cat. No. 14025092), 6.5 mg/mL D-glucose and 0.5% penicillin/streptomycin. Slices were cultured for 5-14 days with full medium changes every 2 days.

### Labeling of CA1 Cells

Dendrites and spines of CA1 pyramidal cells were labelled using a Semliki Forest Virus (SFV)-mediated approach describe in detail elsewhere (Haber et al., 2006). Briefly, SFV driving expression of enhanced green fluorescent protein, targeted to the cell membrane through a farnesylation sequence (EGFPf), was injected into the stratum oriens via pulled glass pipette, broken to a diameter of approximately 50 to 100 μm. Glass pipettes were attached to a Picospritzer III (Parker Hannifin) and SFV was delivered with 10ms pulses at 14 to 18 psi 18 to 20 hours before fixation in 4% formaldehyde/0.1 M PO_4_ ^2-^ for 30 min.

### Confocal Imaging and Spine Analysis

Imaging was performed using an Ultraview Spinning Disc confocal system (Perkin Elmer) attached to a Nikon TE-2000 microscope and a FV1000 laser scanning confocal microscope (Olympus). Z-stacks were acquired from approximately 100µm of CA1 primary apical dendrites, just above the primary dendrite bifurcation. This dendritic subfield is consistently identifiable, fully formed by the period of interest, harbors the highest density of asymmetric synapses, and retains its native connectivity in organotypic slices (Megias et al., 2001; Amaral and Lavenex, 2007). Ten to forty z-stacks were acquired per animal (2-4 animals per experiment, minimum 3 experiments per dataset). Two-dimensional spine counts and geometric measurements of spines were quantified using Reconstruct (Fiala, 2005) and a custom ImageJ macro. 3D spine classification was performed with NeuronStudio (Rodriguez et al., 2008). All spine analysis was performed by an investigator blinded to the experimental condition.

### Western Blot Analysis

For Western blots, 4-6 organotypic slices were lifted from nylon culture inserts with a No. 10 scalpel blade, rinsed in cold PBS and incubated on ice in 100µL of Triton lysis buffer (20 mM Tris pH7.4, 137 mM NaCl, 2mM EDTA, 1% Triton X-100 (TX-100), 0.1% SDS, 10% glycerol, with protease inhibitors and sodium orthovanadate) for 30 min. Lysates were centrifuged at high speed for 10 min and stored at −80°C in sample buffer. Supernatants were warmed to room temperature and run under standard SDS-PAGE conditions. Membranes were immunoblotted with anti-KCC2 1:1000 (N1/12, NeuroMab, CA) and GAPDH 1:300,000 (MAB374, Millipore). KCC2 blots were run immediately after developmental time courses ended to reduce experimentally-induced aggregation of KCC2 oligomers, which we observe to increase with time at −80°C.

### Electrophysiology

Gramicidin perforated patch whole cell recordings were performed similar to previously described (Acton et al., 2012). Briefly, current-voltage (IV) curves were generated by step depolarizing the membrane potential in 10mV increments from ∼-95 to −35mV (Fig 1C) and during each increment GABAergic transmission was elicited via extracellular stimulation in the stratum radiatum. Pipettes had a resistance of 7–12 MΩ and were filled with an internal solution containing 150mM KCl, 10mM HEPES, and 50mM μg/ml gramicidin (pH 7.4, 300 mOsm). We recorded E_GABA_ in current clamp mode. Glutamatergic transmission was inhibited with CNQX

Miniature EPSCs (mEPSCs) were recorded using the whole-cell patch clamp configuration (V_h_ = −70mV), at 30°C, in ACSF containing (in mM): 119 NaCl, 26.2 NaHCO_3_, 11 D-glucose, 2.5 KCl, 1 NaH_2_PO_4_, 2.5 CaCl_2_, 1.3 MgCl_2_, 0.0002 TTX, 0.025 D-APV, 0.05 picrotoxin. Recording pipettes (2-5 MΩ) were filled with (in mM): 122 CsMeSO_4_, 8 NaCl, 10 D-glucose, 1 CaCl_2_, 10 EGTA, 10 HEPES, 0.3 Na_3_GTP, 2 MgATP, pH 7.2. Signals were low-pass filtered at 2kHz, acquired at 10 kHz, and analyzed using Clampfit 10.3 (Molecular Devices).

For cell attached recordings, ACSF and pipette solutions were as described above for mEPSC recordings, but ACSF lacked TTX, D-APV and picrotoxin. Low resistance recording pipettes (1-2 MΩ) were used to form loose patch seals (approximately 100-350 MΩ). Recordings were performed in I=0 mode. GABA was diluted in ACSF to 100 μM and puffed in close proximity to the recorded cell using a glass pipette connected to a Picospritzer III (Parker Hannifin) delivering 10 ms duration air puffs at 14 psi. Electrically-evoked stimulations (1.3 V, 0.5 ms) were delivered by the recording amplifier via the recording pipette. Recorded signals were analyzed using threshold-based detection of spikes in Clampfit 10.3 (Molecular Devices).

Experiments comprised slices from at least 3 separate animals taken from at least 2 litters

### Pharmacology

Pharmacological agents (Tocris unless otherwise noted) were applied to the culture medium during a regular medium change. Gabazine (GBZ) (20µM), bicuculline-methiodide (20µM) and muscimol (10µM) were used to manipulate GABA_A_ transmission. GBZ was washed out by incubating slices in fresh medium for 30 minutes, then washing the top of the slices with equilibrated medium for 1-2 minutes before changing to fresh dishes and medium. Bumetanide (Bume, 10 µM), TrkB-Fc bodies (5mg/mL, R&D Systems) and K252a (200 nM) were added to cultures 30 minutes before adding GBZ.

### Quantitative Reverse Transcriptase

*PCR (qRT-PCR)* Six to eight organotypic slices per sample were lifted from nylon culture inserts with a No. 10 scalpel blade, washed briefly in ice cold PBS and flash frozen in microcentrifuge tubes in a 100% EtOH/dry ice slurry. Total RNA was extracted using the RNeasy Lipid Tissue Kit (Qiagen). cDNA libraries were created using QuantiTect Reverse Transcription Kit (Qiagen). Quantitative PCR was performed using Sybr Green Master Mix (Applied Biosystems Systems) on a StepOne Plus thermocycler (Applied Biosystems). Relative levels of mRNA were calculated using the ΔΔCT method with GAPDH as the internal control. Primer sequences were as follows: GAPDH forward TTG AAG TCG CAG GAG ACA ACC; GAPDH reverse ATG TGT CCG TCG TGG ATC; BDNF forward GTG ACA GTA TTA GCG AGT GGG; BDNF reverse GGG ATT ACA CTT GGT CTC GTA G; Fos forward TCC CCA AAC TTC GAC CAT G; Fos reverse CAT GCT GGA GAA GGA GTC G.

### Immunofluorescence

Slice cultures were fixed as described above, permeabilized for 30 minutes in 1% TritonX 100/PBS, blocked in 10% normal donkey serum (NDS, Jackson Immuno Research)/ 0.2% TX-100/PBS, and incubated with anti-c-Fos antibody (1:5000, Cat. No. 226 003, Synaptic Systems) in 1% NDS/0.2% TX-100/PBS rocking at 4°C for 5 days. Primary antibody solution was washed with 3 rinses in 1% NDS/0.2% TX-100/PBS, followed by secondary antibodies at 1:1000 for 2 hours at room temp. TOPRO-3-iodide (Jackson Immuno Research) was applied at 1:10,000 for 10 minutes in the second of three washes following incubation with secondary antibodies. Quantification of fluorescence intensity with background correction was performed with ImageJ, using mean pixel values in ROIs traced manually around cell bodies.

### Statistics

Data is presented as mean ± SEM. Student t-tests were used except where noted that Mann-Whitney tests were used with datasets with non-normal distribution. Post hoc pairwise comparisons following ANOVA were performed with Tukey’s honestly significant difference (HSD) test. For mean comparisons: *p<0.05, **p<0.01, ***p<0.001. For Kolmogorov-Smirnov tests: ***p<0.0001.

## Acknowledgements

The authors would like to thank Dr. Andrew Greenhalgh and Andy YL Gao for critical review of the manuscript. This work was supported by the Canadian Institutes of Health Research (M.A.W and K.K.M) and Natural Sciences and Engineering Research Council of Canada (K.K.M). C.K.S. was supported through a CGS-M from the CIHR and a Doctoral Award from the FRQS. The authors declare no competing financial interests.

---

**S1 Fig.**
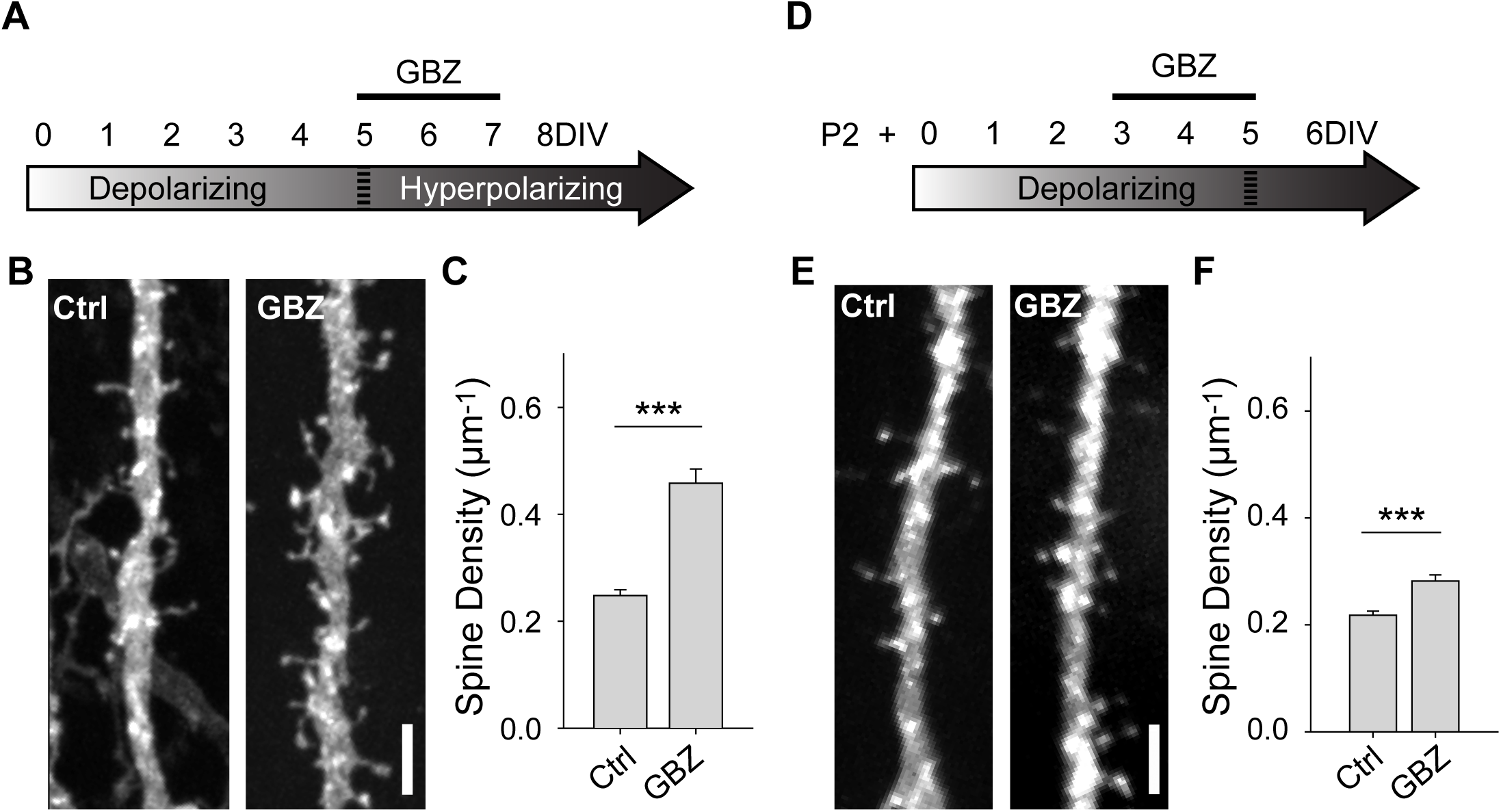
GBZ-induced increase in spines is preserved in absence of antibiotics and in slices cultured from P2 mice. ***A***, Time course of antibiotic-free GBZ treatment. ***B,C,*** Exemplary images and quantification of the spine enhancing effect of GBZ on slices cultures in antibiotic-free culture medium (Ctrl 0.248± 0.0109 μm^-1^, n=198, GBZ 0.458±0.0264 μm^-1^, n=70; N=4; p<0.001, Mann-Whitney). ***D,*** Time course of treatment of slices prepared from P2 pups. ***E,F,*** Exemplary images and quantification of spine enhancing effect of GBZ when applied to slices from P2 pups (Ctrl 0.22 ± 0.008 μm^-1^, n=217, GBZ 0.28 ± 0.01 spines/μm^-1^, n=156; N=3; p<0.001, Mann Whitney).

**S2 Fig.**
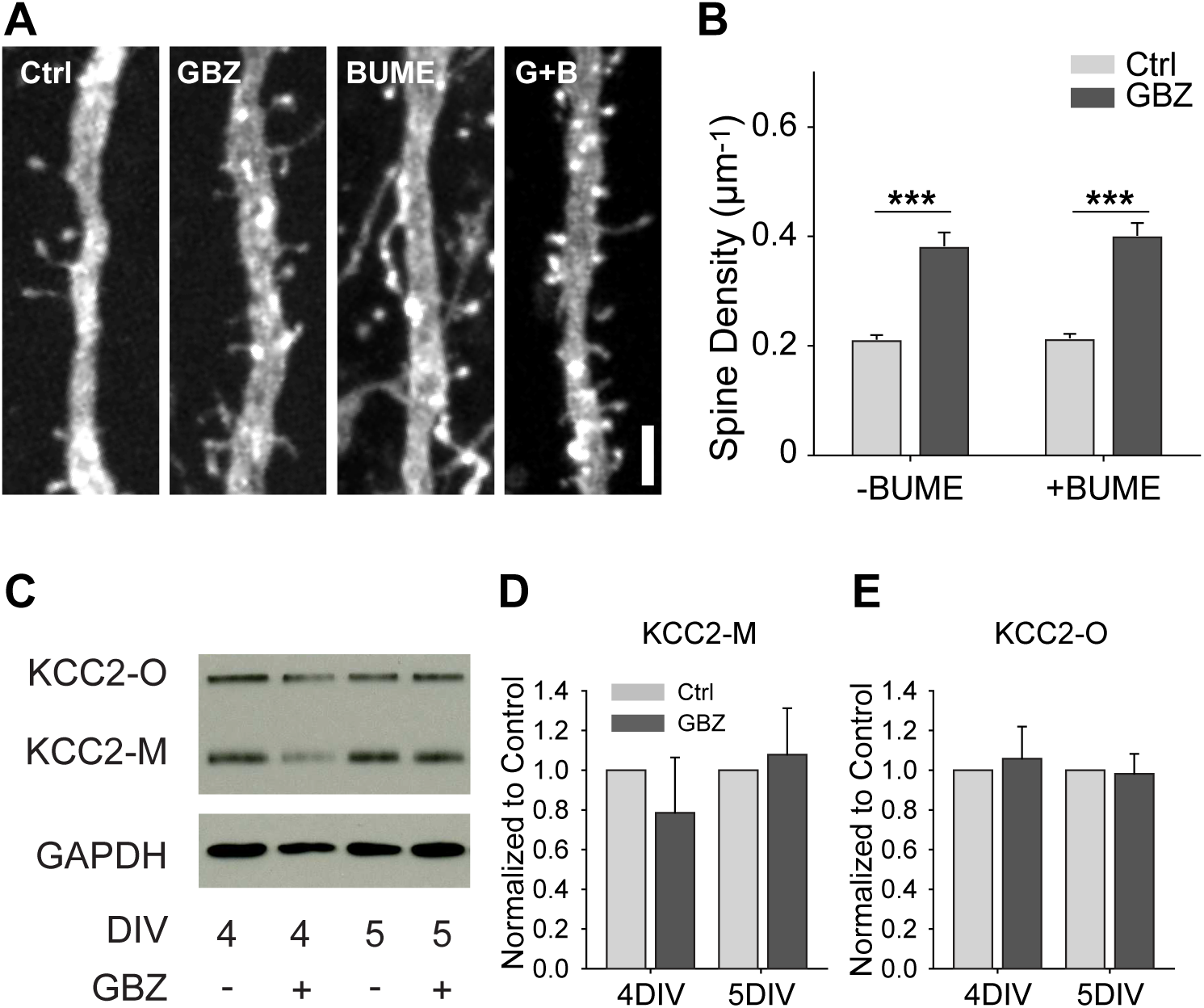
GBZ-induced increase in spines is not reproduced by bumetanide and is not associated with changes in KCC2 expression. ***A,B***, Bumetanide does not increase spine density above control levels, (B, Control 0.21±0.01 um^-1^, n=102; GBZ 0.38±0.02 um^-1^, n=47; BUME 0.21±0.02 um^-1^, n=88_;_ BUME+GBZ 0.40±0.02 um^-1^, n=53_;_ N=3; Two-way ANOVA indicated no significant interaction between GBZ and BUME treatment (p=0.633). Tukey HSD post test indicates significant differences between Ctrl and GBZ in the absence of BUME (p<0.001) and in the presence of BUME (p<0.001)). ***C-E***, Western blot (C) showing no changes in monomeric (D) or oligomeric (E) KCC2 expression following GBZ from 3-4DIV (p=0.52 and 0.77, respectively, One Sample t-Test, n=3) and 3-5 DIV (p=0.76 and 0.87, respectively, One Sample t-Test, n=3). Scale bar 3µm.

**S3 Fig.**
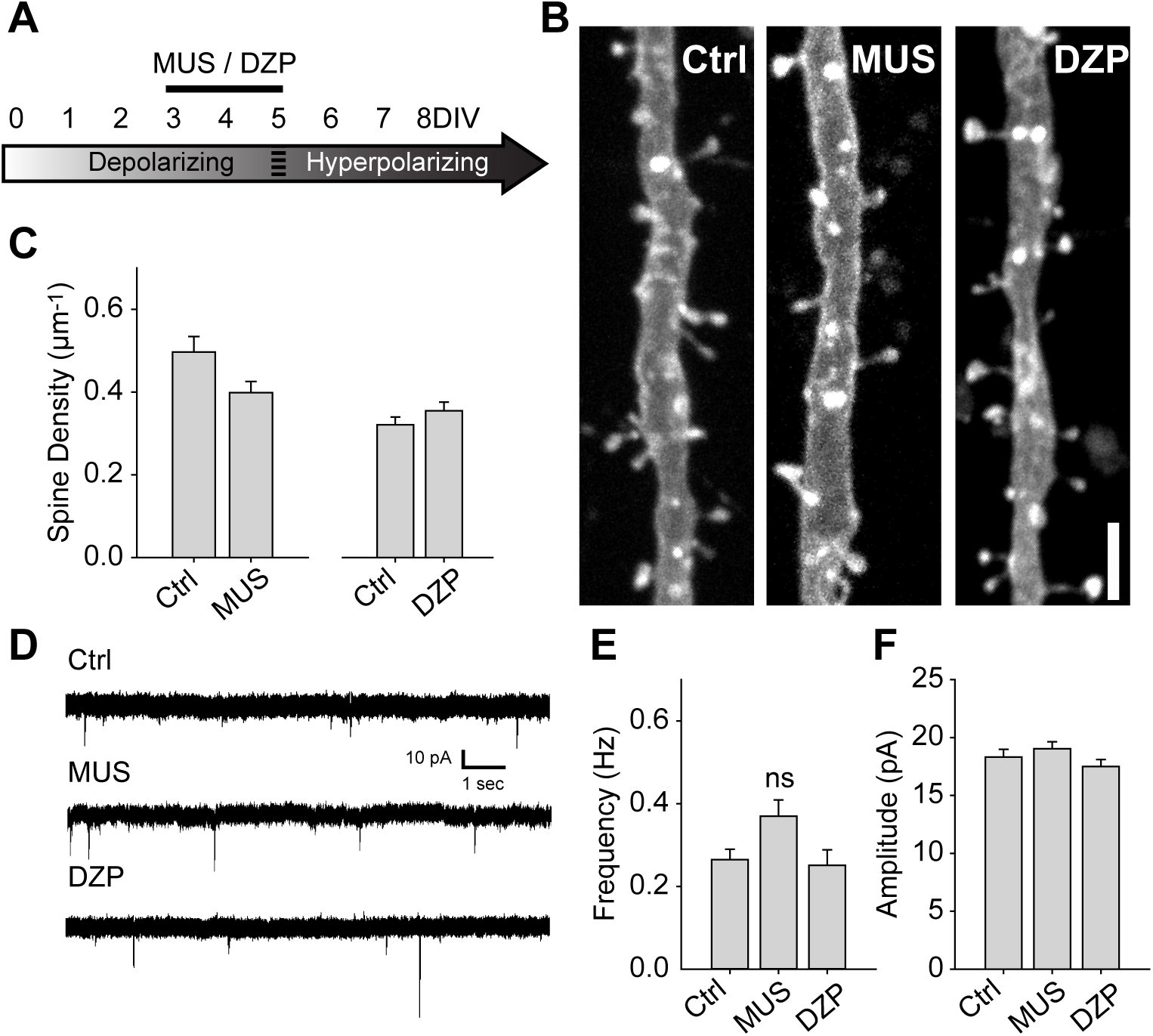
Driving depolarizing GABA_A_ transmission does not decrease glutamatergic synapse numbers. ***A,*** Time course of MUS and DZP treatment. ***B,C***, Spine density after 3-5 DIV MUS treatment (Ctrl 0.50 ± 0.04, n=32; MUS 0.40 ± 0.03, n=39; N=7; p = 0.07) and DZP treatment (Ctrl 0.321 ± 0.02, n=116; DZP 0.36 ± 0.02, n=88; N=6; p = 0.11, Mann-Whitney). ***D,*** Representative traces of mEPSCs following 3-5DIV treatment with MUS or DZP. ***E,*** mEPSC frequency summary plot (Ctrl: 0.27 ± 0.02 Hz, n=9; MUS: 0.37 ± 0.04 Hz, n=8; DZP: 0.25 ± 0.04 Hz, n=8; One way ANOVA p=0.046; Ctrl vs MUS, p=0.09; Ctrl vs DZP, p=0.9). ***F,*** mEPSC amplitude summary plot (Ctrl: 18.3 ± 0.7 pA, n=9; MUS: 19.0 ± 0.6 pA, n=8; DZP: 17.5 ± 0.6 pA, n=8; One Way ANOVA, p=0.263)

**S4 Fig.**
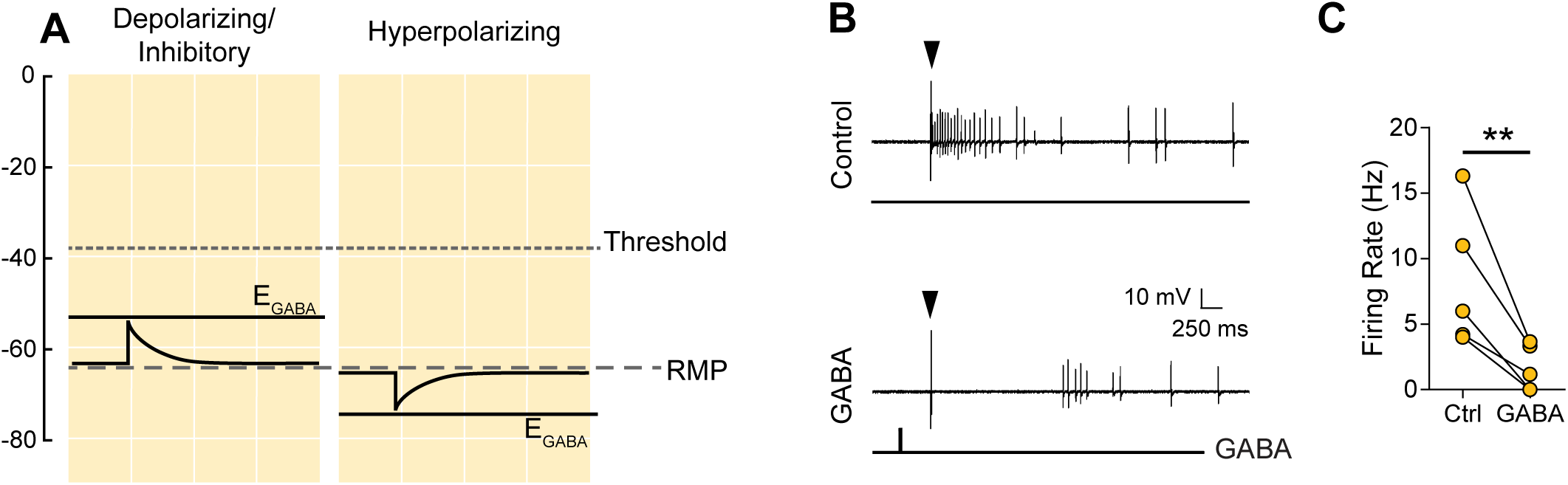
Shunting GABA transmission inhibits electrically evoked firing in CA1 neurons at 3DIV. ***A,*** Schematic demonstrating the likely shunting and hence inhibitory nature of GABA_A_ transmission due to the relative values of AP Threshold>E_GABA_>RMP. The scale in A aligns with that of Fig 1 ***E, F*** and ***G*** such that the threshold, RMP and E_GABA_ values are represented accurately relative to each other. ***B,*** Sample traces from the same cell demonstrating that activity could be evoked electrically (Control) and that puffed GABA inhibited electrically evoked activity (GABA). The arrow above the traces denotes the timing of electrical stimulation. ***C,*** Summary plots of electrically evoked activity in the absence and presence of puffed GABA.

**S5 Fig.**
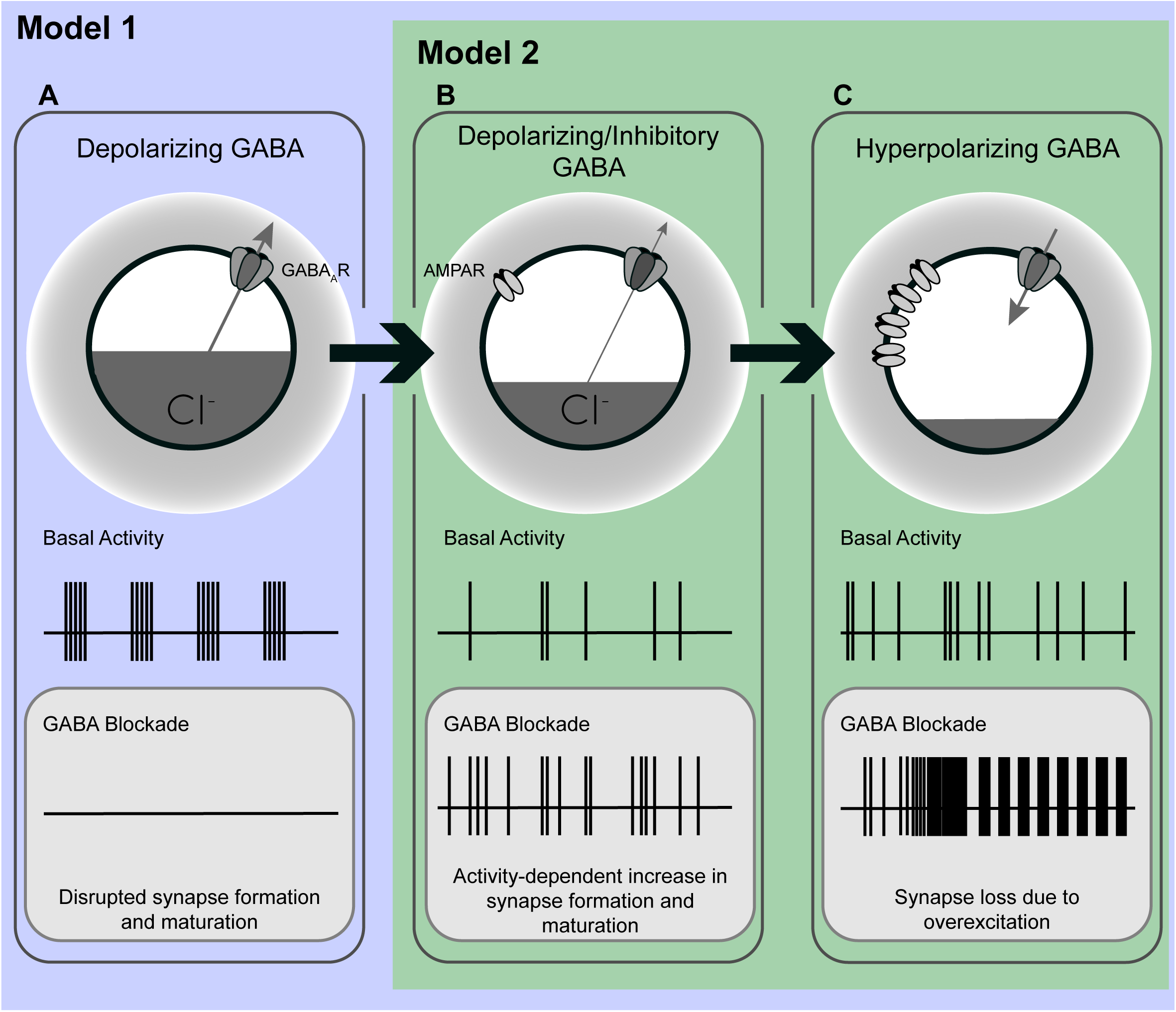
A model of the possible roles of GABA_A_ transmission in glutamatergic synapse formation as chloride homeostasis matures. ***A,*** Work performed in acute slices suggests that depolarizing GABA_A_ transmission provides the initial excitatory drive required for activity- and calcium-dependent formation and maturation of glutamatergic synapses. Blocking GABA_A_ transmission at this stage eliminated GDPs. For the sake of simplicity we have depicted that GABA_A_ blockade would eliminate GDPs and silence network activity at this stage, however it should be noted that in acute slices, blocking GABA_A_ transmission at this point has been shown to decrease circuit activity in immature acute hippocampal slices as depicted (Ben-Ari et al., 1989; Garaschuk et al., 1998; Mohajerani and Cherubini, 2005), but has also been shown to induce interictal discharges (Khazipov et al., 1997; Khalilov et al., 1999; Lamsa et al., 2000) or paroxysmal activity (Wells et al., 2000). These latter effects may be due to an overarching inhibitory role for GABA during development. ***B,*** Our work suggests a possible transition state wherein blocking GABA_A_ transmission alleviates a depolarizing but inhibitory restraint on circuit activity, allowing for activity dependant formation of glutamatergic synapses. Such a transition state would likely rely on a still underdeveloped glutamatergic system that is not yet capable of pathological levels of overexcitation. Importantly, recent *in vivo* work suggests that GABA may be inhibit circuit activity throughout postnatal development, indicating that blocking GABA_A_ transmission might enhance glutamatergic synapse formation from birth until GABA becomes fully hyperpolarizing. (Although the basal activity here is depicted as uncoordinated to clearly differentiate A from B, the activity pattern in this transition state, as well as in C, may very well be composed of ENOs.) ***C,*** When E_Cl_ and the glutamatergic system are mature, blocking hyperpolarizing GABA_A_ transmission causes overexcitation and loss of glutamatergic synapses.

